# Mapping excitatory synaptic plasticity evoked by single-dose psilocybin in mice

**DOI:** 10.64898/2026.07.31.741703

**Authors:** Ziming Li, Crystal Weber, Francesca Sellitti, Linda D. Simmler

## Abstract

A single dose of psilocybin can induce long-lasting antidepressant effects. The neurobiological mechanisms underlying such sustained antidepressant effect remain insufficiently understood, particularly at the level of synaptic function and drug-target specificity. Here, we aimed to delineate single-dose psilocybin-induced excitatory synaptic plasticity. Synaptic plasticity was assessed by whole-cell patch-clamp recording of excitatory synaptic transmission 24 h after treating mice with single-dose psilocybin. We correlated the recordings with transcriptomics data and used a conditional single-vector CRISPR/SaCas9-dependent knock-out strategy to validate the role of the 5-HT_2A_ receptor. Psilocybin selectively increased the frequency of miniature excitatory postsynaptic currents in specific cortical sub-regions and in the amygdala. Frequency correlated with the expression levels of psilocin-targeted serotonin receptors, when expression heterogeneity between cortical subregions and along the anterior-posterior axis was accounted for. Post-synaptic *Htr2a* knock-out in the insular/orbitofrontal cortex precluded psilocybin-induced 24-h plasticity. These findings demonstrate that lasting psilocybin-induced effects on excitatory synaptic transmission manifest with brain-region specificity, likely reflecting a functional consequence of synapse formation. This work establishes a foundation for a circuit-specific, mechanistic understanding of functional aspects of psilocybin-induced neuroplasticity.

## Introduction

Psilocybin is a serotonergic psychedelic that has drawn widespread attention in recent years for its potential therapeutic value in treating depression^1–4^. With clinical studies showing that one or few administrations can produce rapid and sustained antidepressant effects, these findings have intensified interest in the neural mechanisms that underly its therapeutic actions outlasting acute drug effects. Among the leading hypotheses is that psilocybin acts as a plasticity-inducing drug, triggering enduring forms of structural and functional plasticity in brain circuits implicated in affective regulation^5–7^.

Preclinical studies have shown that a single dose of psilocybin can induce rapid and persistent dendritic spine growth in frontal cortical neurons^5^ and alter molecular programs linked to synaptic remodeling^8,9^. However, current research on psilocybin and related psychedelics remains heavily focused on a limited number of brain regions, predominantly the medial prefrontal cortex^10,11^ and the anterior cingulate cortex^5,12,13^ as well as structural rather than functional consequences of neuronal plasticity. There is a lack of systematic studies examining whether psilocybin-induced changes in excitatory synaptic transmission are specific or generalizable across brain regions, including sub-regions of the frontal cortex and non-cortical areas. This gap limits our understanding of functional aspects of psychedelic-induced neuroplasticity and obstructs efforts to dissect circuit- and cell-type specific effects in the context of brain-wide drug.

Psilocybin exerts its effects through multiple serotonin (5-HT) receptor subtypes, as its active metabolite psilocin activates potently 5-HT_2A_, 5-HT_2C_, and 5-HT_1A_ receptors^14,15^. The 5-HT_2A_ receptor is widely considered a critical mediator of psychedelics’ canonical neural and behavioral effects^11,13,16,17^. Yet, whether heterogeneity of *Htr2a* expression across cortical regions predicts regional differences in psilocybin-induced synaptic plasticity, has not been systematically assessed. Recent large-scale MERFISH-based mouse brain atlases^18,19^ provide an unprecedented opportunity to address this question, when combined with spatially resolved neurophysiological data.

Here, we aimed to determine lasting psilocybin-induced excitatory synaptic plasticity across brain regions and identify contributions of drug-targeted receptors. We generated a multi-regions whole-cell patch-clamp data set for miniature excitatory postsynaptic currents (mEPSCs) recorded 24 h after single-dose psilocybin. We report that psilocybin-induced changes in excitatory synaptic transmission are spatially organized and related to 5-HT receptor expression levels, with the 5-HT_2A_ receptor being necessary for the induction of psilocybin-induced increase of mEPSC frequency.

## Results

### Psilocybin consistently induced transient increase in head-twitching

In order to validate the efficacy of psilocybin injections, we quantified head-twitch response to a single intraperitoneal (i.p.) injection of 1 mg/kg psilocybin in mice (Figure 1a). Head-twitching occasionally occurred at baseline, and psilocybin transiently increased head-twitching (Figure 1b) in each of the injected animals (Figure 1c), supporting the validity of the psilocybin treatment for patch-clamp experiments.

**Figure 1:**
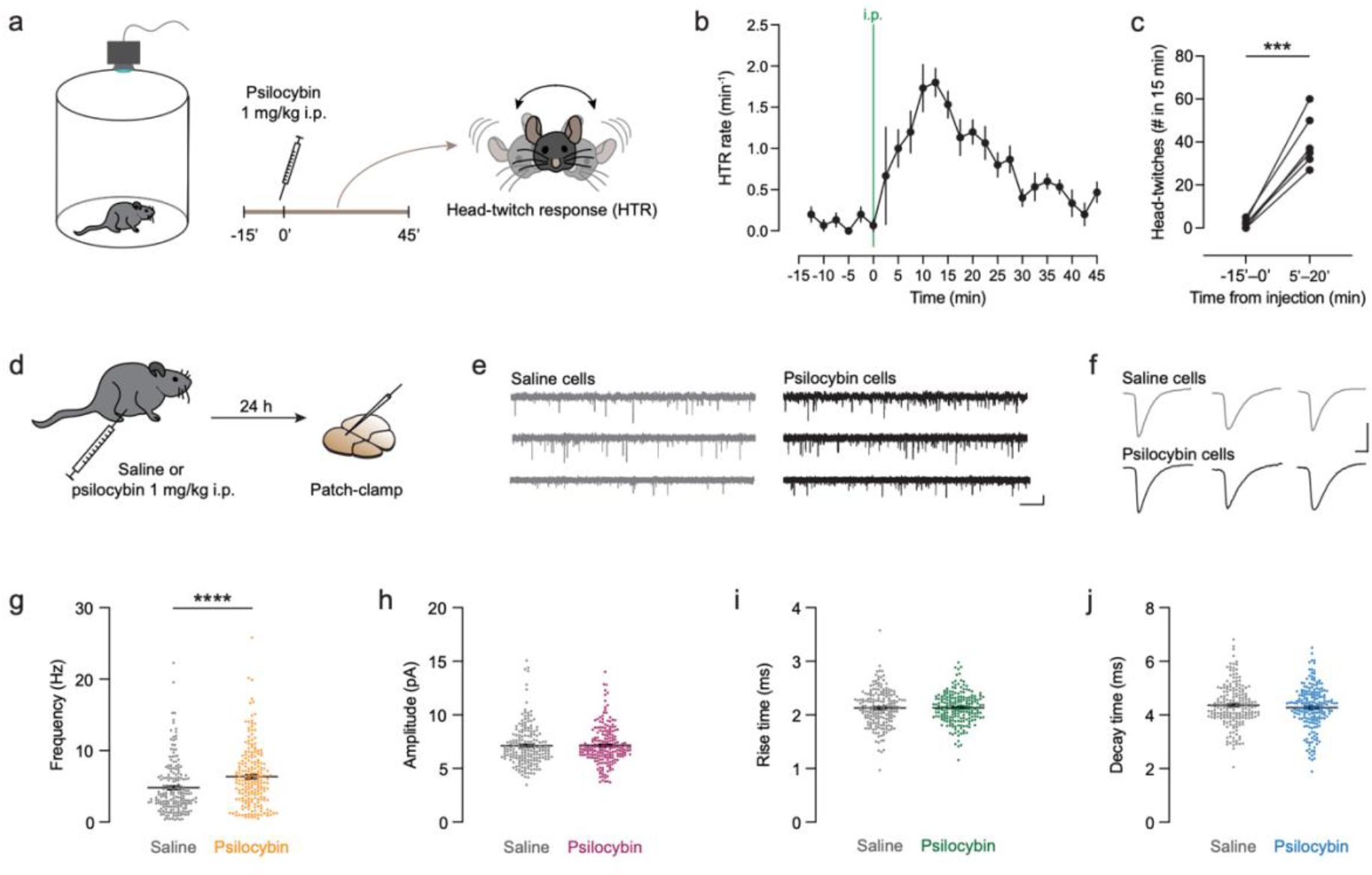
Synaptic effects of single-dose psilocybin 24 h after injection. **a)** Experimental design for behavioral confirmation of psilocybin efficacy. **b)** Time-dependent increase in head-twitch response (HTR) rate after i.p. injection of 1 mg/kg psilocybin. N=6 mice. **c)** Number of head-twitches in 15 min increased in all mice compared to baseline. ***p=0.0004, paired t-test, N=6. **d)** Experimental design for assessing synaptic effects at 24 h after treatment with psilocybin 1 mg/kg or saline in cortical areas, hippocampus and amygdala. **e)** Example traces of recordings of six representative cells from saline and psilocybin-treated mice. Scale bar is 1 s, 5 pA. **f)** Averaged events of six representative cells. Scale bar is 5 ms, 5 pA. **g)** Frequency of mEPSC events from psilocybin-treated mice was significantly elevated compared to saline. ****p<0.0001, Mann-Whitney comparison. **h–j)** No significant differences in mEPSC amplitudes **(h)**, rise time **(i)**, and decay time **(j)** between saline-and psilocybin-treated mice. Data in b, g–j are mean±SEM, individual data points in g–j are recorded cells, N=192 cells from 34 mice (saline); 200 cells from 35 mice (psilocybin).

### Increased mEPSC frequency 24 h after a single psilocybin injection

To characterize lasting excitatory synaptic plasticity from a single injection of psilocybin, we injected mice 24 h before sacrifice for patch-clamping with psilocybin (1 mg/kg) or saline (Figure 1d). The psilocybin dose was chosen based on previous research with mice^5,20,21^. For patch-clamp recordings, we targeted layer 5/6 cortical cells in different cortical areas. Furthermore, we included cells in the hippocampus and the amygdala. We assessed frequency, amplitudes, rise time and decay time of mEPSCs from a total of 392 cells (example traces in Figure 1e, f). Compared to saline-treated mice, mEPSC frequency in cells from psilocybin-treated mice was significantly elevated (Figure 1g). Psilocybin increased median frequency by 52% (3.8 Hz for saline vs. 5.8 Hz for psilocybin). Amplitude, rise- and decay time did not differ between cells from psilocybin compared to saline-treated mice (Figure 1h–j). The increase of mEPSC frequency observed 24 h after a single dose reflects long-lasting neuronal changes, as the acute effect of psilocybin decreases quickly in mice, evidenced by the head-twitch response returning close to baseline by 45 min after injection (Figure 1c). Increased mEPSC frequency reflects either a psilocybin effect on a functional through increased presynaptic release probability^22^, or on a structural level through more functional synapses^23,24^. The net effect of both possibilities is stronger synaptic signaling.

### Spatial mapping of mEPSC frequency identifies brain regions with synaptic plasticity

Spatial resolution allows to infer findings flexibly to brain areas and subregions, allowing interpretation by brain area function. We therefore kept track of recorded cells during patch-clamping using a self-made code “Mark where you patched”, through which the patch-clamper could easily register the approximated cell location in an interface based on the Allen Mouse Brain Common Coordinate Framework (CCFv3)^25^. To illustrate the mEPSC frequency results in a spatial map, we normalized mEPSC frequencies to saline within brain areas as categorized by the brain atlas and displayed normalized frequencies for all recorded cells in the Allen brain framework (Figure 2a). The mapping illustrates the even distribution of saline and psilocybin cells and implies a predominant effect in anterior cortical areas.

**Figure 2:**
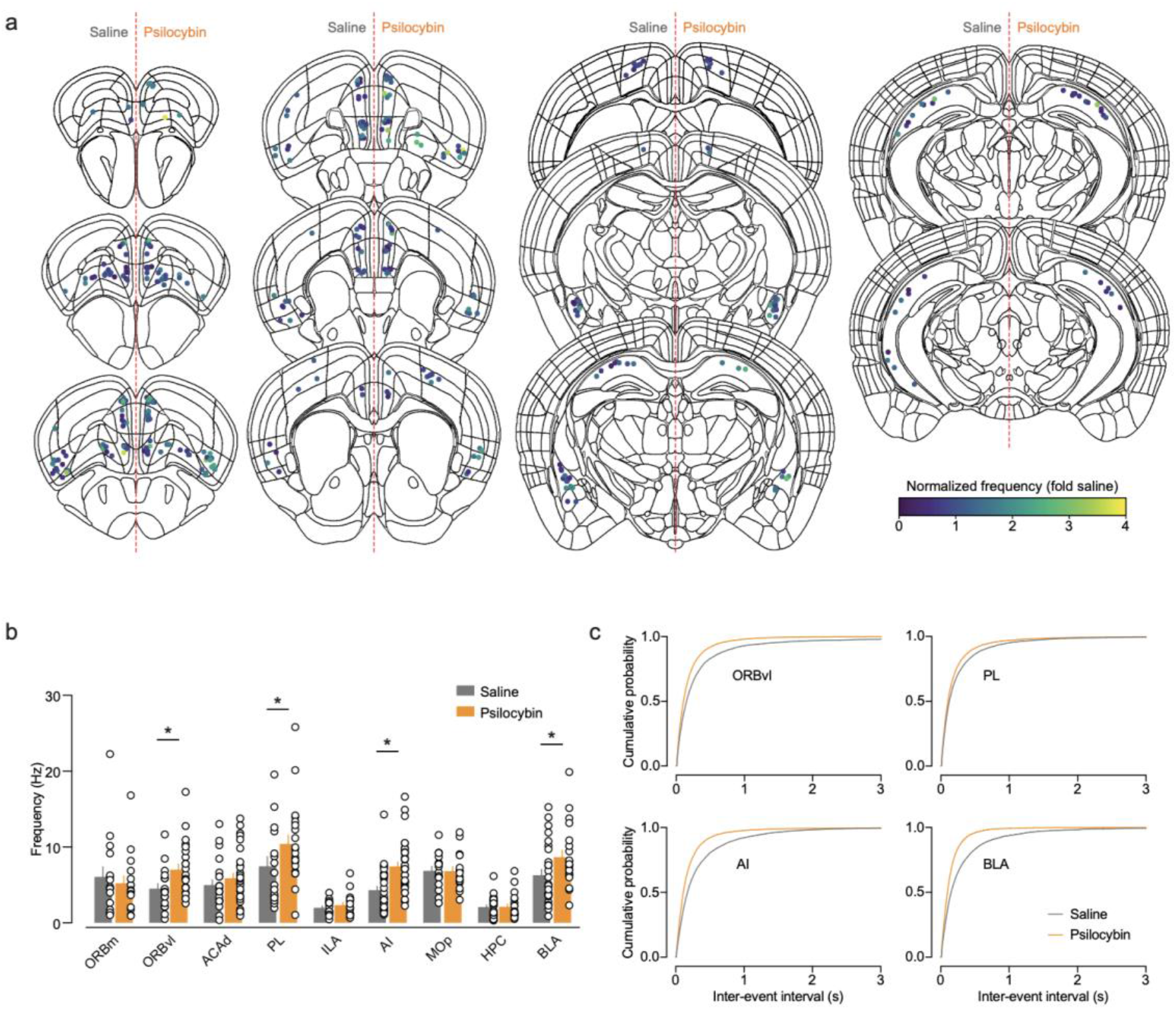
Spatial resolution of psilocybin-induced synaptic effects. **a)** Location of recorded cells from Figure 1 plotted in the Allen brain mouse atlas and colored by individual mEPSC frequencies normalized to saline within brain areas (as defined in b). Cells from saline-treated mice in left hemisphere, from psilocybin-treated mice in right hemisphere. **b)** mEPSC frequency resolved by targeted brain area. Psilocybin significantly elevated frequency in the ORBvl, PL, AI and BLA. Significant interaction (p=0.041) in 2-way ANOVA, *p<0.05 in multiple comparison with FDR correction. N(each condition)=16–35 cells from 5–8 mice (Table S1 reporting exact sample size). Data are mean±SEM with cells as individual data points. **c)** Mean cumulative curves for inter-event intervals from ORBvl, PL, AI and BLA. ORBm, medial orbital area; ORBvl, ventrolateral orbital area; ACAd dorsal anterior cingulate area; PL, prelimbic area; ILA, infralimbic area; AI, agranular insular area; MOp, primary motor area; HPC, hippocampus CA1; BLA, basolateral amygdala.

Then, we run a two-way ANOVA on mEPSC frequencies segregated by brain areas and confirmed a treatment effect for mEPSC frequency (Figure 2b). Post-hoc testing identified significant frequency elevation in cells from psilocybin-treated mice in the ORBvl, PL, AI and BLA, but not in the ORBm, ACAd, ILA, MOp and HPC (Figure 2c). Selectivity of drug effects to these brain areas is also reflected in the cumulative curves of inter-event intervals (Figure 2c, Figure S1). Of note, mean frequencies of the saline conditions were different between brain regions (Figure 2a), which was evident by a significant main effect for brain regions. Amplitudes of the mEPSCs were not statistically different between saline- and psilocybin-treated mice when separating the data set by brain area (Figure S2a), although cumulative curves of single areas implied isolated drug effects (Figure S2b).

### Opposing changes in decay time kinetics in the ACAd and BLA

Segregation of the mEPSC decay time by brain area resulted in a significant 2-way ANOVA interaction (Figure 3a). With psilocybin-treatment, decay time was decreased in the ACAd and increased in the BLA, compared to the saline condition (Figure 3a). Decay and rise time can positively correlate^26^, but are independent variables^27^. We therefore plotted decay times of ACAd and BLA cells against their rise time. In both brain areas, decay and rise times strongly correlated (Figure 3b, c). Accordingly, rise time was decreased in the psilocybin than in the saline condition for cells in the ACAd (Figure 3b). These findings show that psilocybin-induced synaptic plasticity can involve changes in mEPSC kinetics, indicative of altered AMPA receptor subunit composition from psilocybin treatment^28^ and dendritic filtering^26^, but altered kinetics were present only in two of the tested brain areas and in opposing directions.

**Figure 3:**
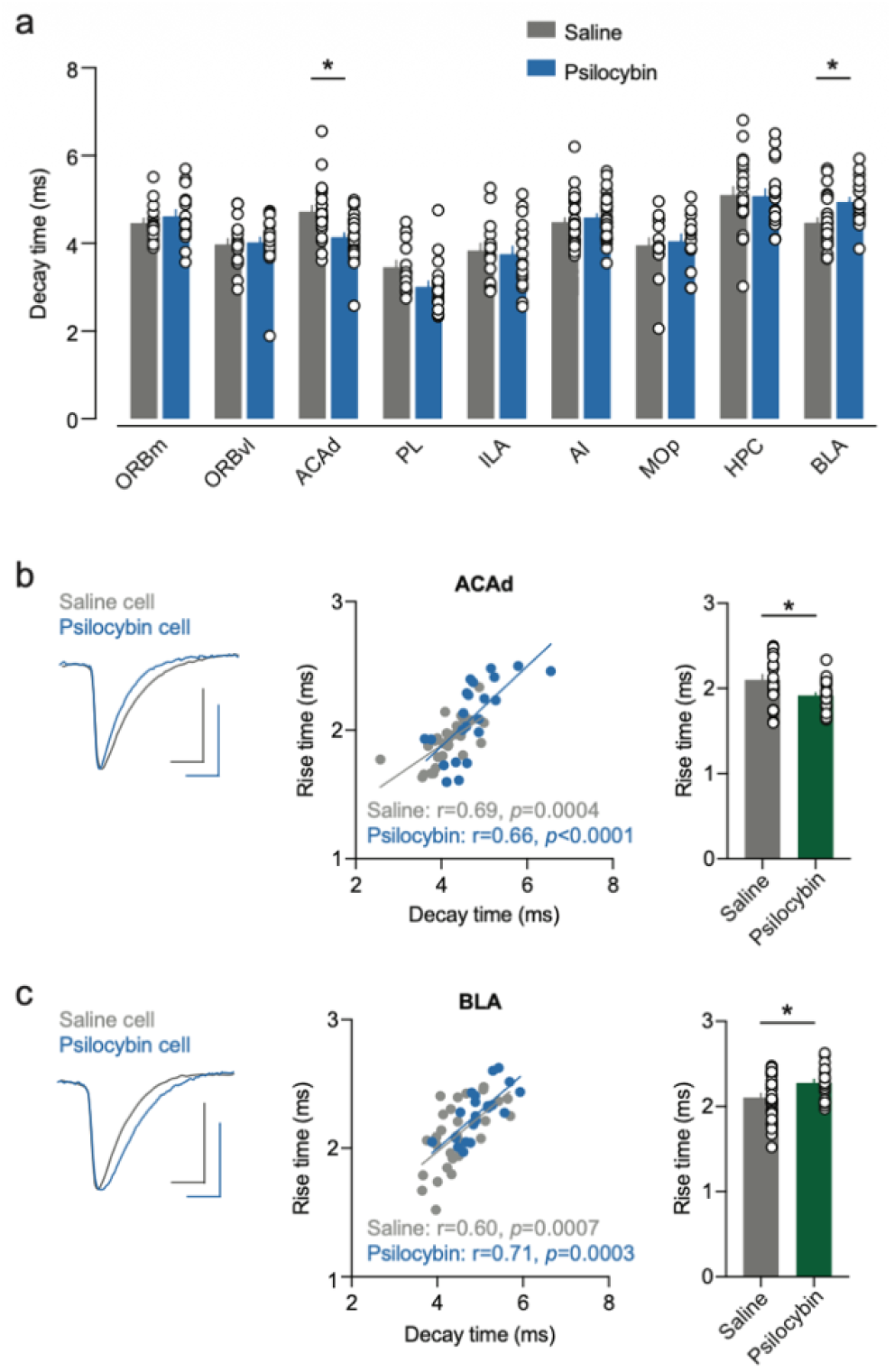
Effects of psilocybin on mEPSC kinetics. **a)** mEPSC decay time from Figure 1 resolved by targeted brain area. Psilocybin significantly changed decay time in the ACAd and BLA. Significant interaction (p=0.0022) in 2-way ANOVA, *p<0.05 in multiple comparison with FDR correction. N(each condition)=16–35 cells from 5–8 mice (Table S1 reporting exact sample size). **b)** Left: Representative averaged events of a saline- and a psilocybin cell in the ACAd. Middle: Rise time significantly correlated with decay time in the ACAd. Pearson correlation. Right: Rise time was significantly decreased in psilocybin-treated mice. *p=0.012, Welch’s corrected unpaired t-test. **c)** Left: Representative averaged events of a saline- and a psilocybin cell in the BLA. Middle: Rise time correlated with decay time in the BLA. Pearson correlation. Right: Rise time was significantly increased in psilocybin-treated mice. *p=0.010, Welch’s corrected unpaired t-test. Summary data are mean±SEM, dots are individual cells. Scale bars are 5 ms, 5 pA. ORBm, medial orbital area; ORBvl, ventrolateral orbital area; ACAd dorsal anterior cingulate area; PL, prelimbic area; ILA, infralimbic area; AI, agranular insular area; MOp, primary motor area; HPC, hippocampus CA1; BLA, basolateral amygdala.

### 5-HT2A receptor expression correlated with mEPSC frequency

The spatial mapping of our mEPSC results allowed to correlate these data with open-source data with the same spatial information. To relate our data to expression levels of psilocin’s main pharmacological targets, we first plotted gene expression levels from a transcriptomics data set imputed to spatial location^19^, including only gene expression from glutamatergic cells. Expression of *Htr2a* was relatively consistent across the selected cortical areas (Figure 4a), yet differences are apparent, such as relatively low levels in the ILA and high levels in insular areas (AId, AIp, AIv, GU). In contrast, the hippocampal CA1 area and the basolateral amygdala expressed *Htr2a* to a lesser extent than the cortical areas, whereas their *Htr2c* expression was more prominent, particularly for the BLA (Figure 4a). For *Htr1a* expression, CA1 stood out as area with most prominent expression (Figure 4a). We plotted glutamatergic gene expression in the anterior-posterior axis for the cortical areas (Figure 4b). Similar to our observation with mEPSC frequency plotted in the brain atlas framework in Figure 2, we observed an anterior to posterior gradient for *Htr2a* expression across and within cortical brain regions (Figure 4b). This motivated us to correlate mEPSC frequencies in cortical areas with gene expression levels of psilocin’s major targets *Htr2a*, *Htr2c* and *Htr1a*, with gene expression from glutamatergic cells in 0.5 mm anterior-posterior bins from bregma (same as in Figure 4b). Interestingly, in the psilocybin condition, all three target genes correlated significantly, although weakly, with mEPSC frequencies (Figure 4c). In contrast, the no correlation was evident for the vesicular glutamate transporter gene *Slc17a7*, which served as control (Figure 4c). In the saline condition, we also found a correlation between frequency and gene expression for *Htr2a*, but not for *Htr2c* and *Htr1a* (Figure 4c). By comparing mapped patch-clamp mEPSC data to spatial transcriptomics, we found that expression levels of the drug-targeted 5-HT receptors are associated with psilocybin-induced increase in mEPSC frequency, yet with small effect size of the correlations.

**Figure 4:**
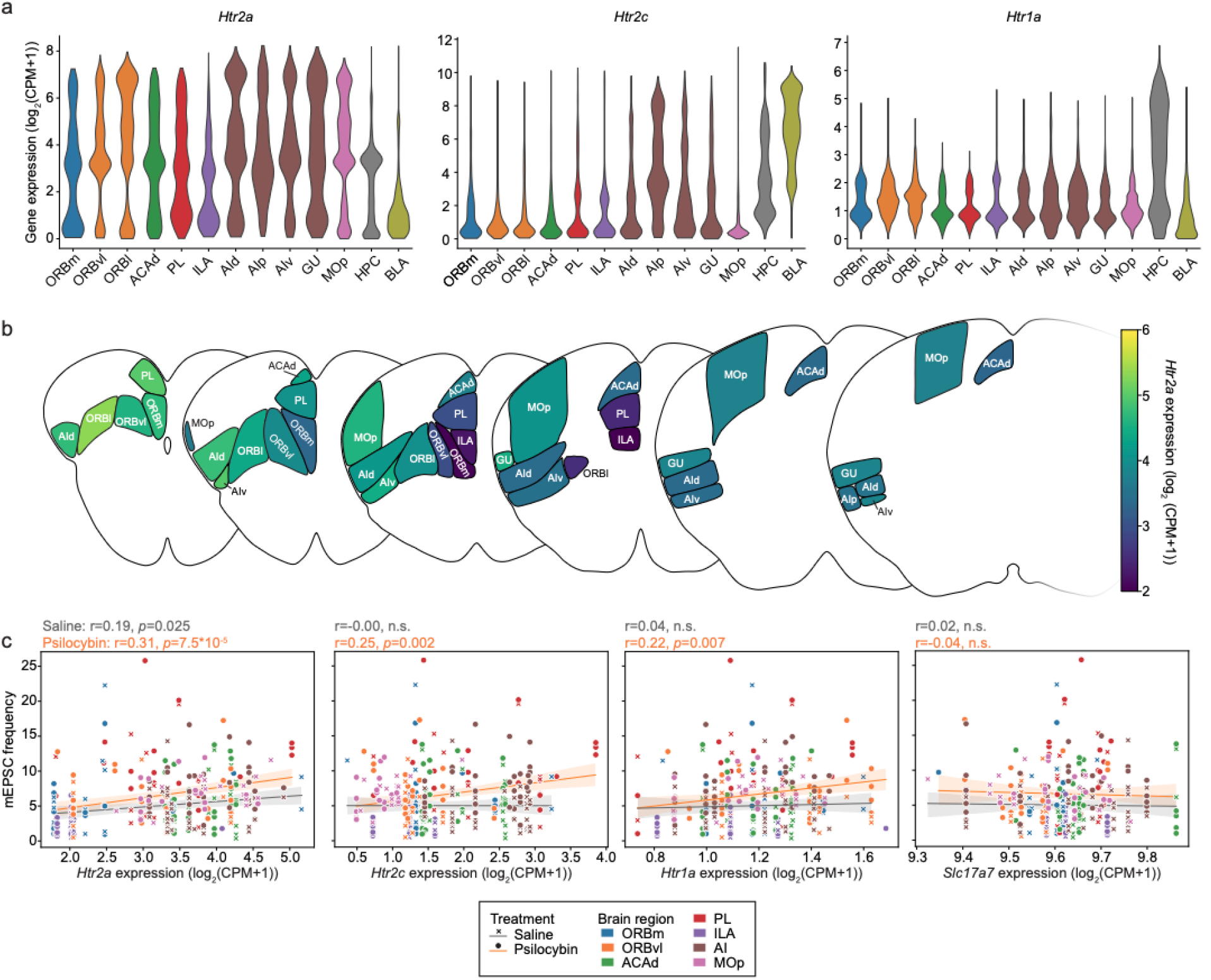
Correlation of mEPSC frequency with spatially resolved 5-HT receptor expression levels. **a)** Expression levels of Htr2a (left), Htr2c (middle), and Htr1a (right) in glutamatergic cells in brain regions with mEPSC recordings. Expression levels were extracted from the data set by Yao et al.^19^. Regions with identical colors were combined in for analysis in c. **b)** Htr2a expression as in a, but expression resolved by anterior-posterior location. **c)** Levels of gene expression at respective anterior-posterior location correlated with mEPSC frequency resolved by brain area as in Figure 2. Linear regression within treatment with 95% CI. ORBm, medial orbital area; ORBvl, ventrolateral orbital area; ORBl, lateral orbital area; ACAd dorsal anterior cingulate area; PL, prelimbic area; ILA, infralimbic area; AId, dorsal agranular insular area; AIp, posterior agranular insular area; AIv, ventral agranular insular area; GU, gustatory areas; Mop, primary motor area; HPC, hippocampus CA1; BLA, basolateral amygdala.

### Area-specific Htr2a knock-out precluded psilocybin-induced 24-h plasticity locally

For more insights into the relationship between drug-targeted receptor expression and synaptic plasticity, we set out to determine to what extent the 5-HT_2A_ receptor, the main and shared target of serotonergic psychedelic compounds, contributed to the psilocybin-induced excitatory synaptic plasticity which we described above. We therefore employed a Cre-dependent, single vector CRISPR/SaCas9-dependent knock-out strategy^29^ to remove *Htr2a* in neurons located in the insular area/ventrolateral orbital area (Figure 5a) where *Htr2a* expression and psilocybin-induced frequency increase were prominent (Figures 2b, 4a). *Rosa26* knock-out (Rosa26) served as control^29^. Four weeks after stereotactic surgeries, mice were treated with saline or psilocybin and mEPSCs patch-clamp recordings were performed in fluorescent cells (Figure 5b for representative biocytin-filled patched cells). Location of patched cells were tracked with our newly developed “Mark where you patched” interactive viewer and plotted for visualization (Figure 5c). While psilocybin-induced increase in mEPSC frequency from Figure 2b was reproduced in the Rosa26 control condition, psilocybin treatment did not increase mean mEPSC frequency when *Htr2a* was locally knocked out (Figure 5d). When plotting inter-event intervals as cumulative curves, a small change in the larger intervals was still evident in the *Htr2a*-KO psilocybin condition compared to saline controls, but the change was clearly smaller than in the Rosa26 control data (Figure 5e). Comparable to our findings in mice without viral transduction, no drug- or genotype effects were observed on amplitude (Figure 5f), decay time (Figure 5g) and rise time (Figure 5h). These findings indicate that postsynaptic 5-HT_2A_ receptors play a major role in the induction of psilocybin-induced mEPSC frequency changes and points to the importance of post-synaptic expression of the drug target.

**Figure 5:**
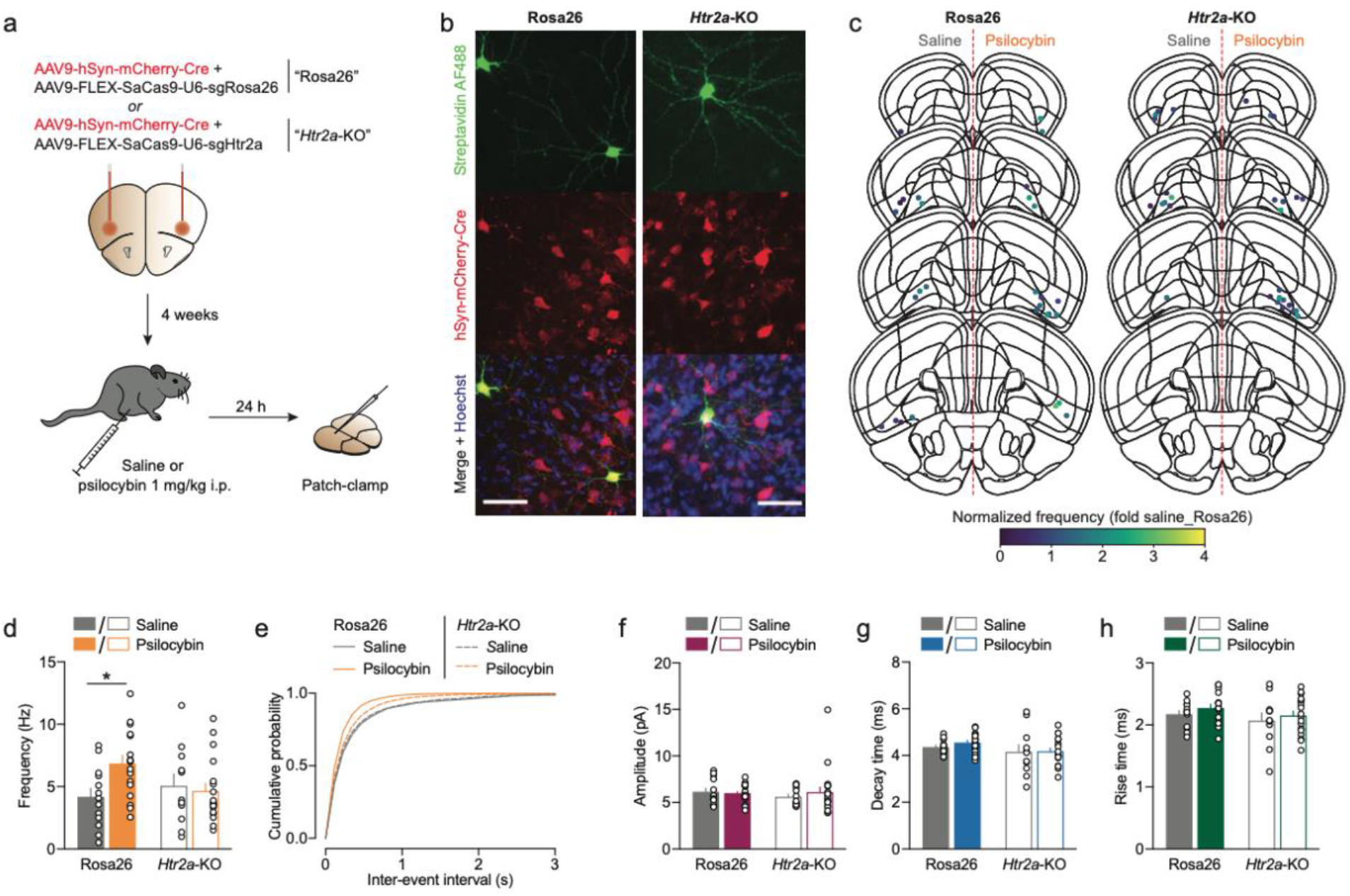
Lack of psilocybin-induced frequency increase with local knock-out of Htr2a. **a)** Experimental design. Virus constructs for local KO of Htr2a and control were injected targeting the insular area/ventrolateral orbital area. After 4 weeks, mice were treated and patch-clamping performed 24 h thereafter. Fluorescent-positive neurons were recorded. **b)** Representative images of biocytin-filled patched cells for the Rosa26 control and the Htr2a-KO conditions (green, streptavidin Alexa Fluor 488 (AF488); red, mCherry expression from virus delivering Cre; green/red/blue (Hoechst-stained nuclei) merged). **c)** Location of recorded cells plotted in the Allen brain mouse atlas and colored by individual mEPSC frequencies normalized to in the Rosa26 saline condition (data from d). Cells from saline-treated mice in left hemispheres, from psilocybin-treated mice in right hemispheres. **d)** Significantly increased frequency of mEPSCs with psilocybin treatment in Rosa26 controls but not in cells with Htr2a KO. Significant interaction (p=0.038) in 2-way ANOVA, *p<0.05 with Sidak post-hoc test. **e)** Cumulative probability curves for data from d. **f)** Same recordings as in d, but amplitude plotted. No significant differences. **g)** Same recordings as in d, but decay time plotted. No significant differences. **h)** Same recordings as in d, but rise time plotted. No significant differences. Data in d–h are mean±SEM (except for e where data are mean only), individual data points are recorded cells, N=13 cells from 3 mice (Rosa26 saline); 17 cells from 3 mice (Rosa26 psilocybin), 11 cells from 3 mice (Htr2a-KO saline), 18 cells from 4 mice (Htr2a-KO psilocybin).

## Discussion

Here, we demonstrate that a single dose of psilocybin induces robust and spatially confined alterations of excitatory synaptic function, a drug effect involving 5-HT_2A_-receptors and expressed the day after drug administration. With this, we present the to date largest data set on psychedelics-induced synaptic plasticity across brain areas. Psilocybin increased mEPSC frequency, while mEPSC amplitudes were unchanged. The frequency effect was not homogenously distributed across investigated brain areas, but confined to the ORBvl, PL, AI, BLA. Transient kinetics were selectively modulated by psilocybin in the ACAd and BLA with opposing effects. The involvement of drug-targeted 5-HT receptors was evident by positive correlations between expression levels and mEPSC frequency, as well as the necessity of the 5-HT_2A_ receptor for drug-mediated mEPSC frequency changes. Together, these findings support a model in which psilocybin induces long-lasting, 5-HT receptor-mediated and brain region-specific functional changes of excitatory synaptic inputs that might reflect clinically relevant circuit reorganization.

Complexity of psychedelic-induced synaptic plasticity has been highlighted in a systematic review summarizing electrophysiological studies with mostly ex vivo application of psychedelic drugs, concluding that psychedelics modulate excitation and inhibition in a cell-type- and region-specific, biphasic, and dose-dependent manner rather than uniformly raising cortical excitability^30^. Here, the increase in mEPSC frequency provides direct functional evidence that excitatory synaptic input to the recorded neurons was enhanced 24 h after in vivo psilocybin treatment. This reflects neuroplasticity persistent beyond acute drug effect, which, as measured here by head-twitch-response, subsided within 45 min. This persistence is consistent with long-lasting structural remodeling of frontal-cortical dendritic spines reported after a single psilocybin dose^5,13^ and the broader capacity of serotonergic psychedelics to induced lasting structural and functional plasticity^10^. The interpretation of mEPSC parameters is based on the understanding that recorded events are action-potential-independent quantal release of neurotransmitters. The significant increase in mEPSC frequency shows that recorded neurons received more frequently detectable AMPAR-mediated excitatory events a day after psilocybin treatment, compared to saline treatment. Our experiments did not determine to what extent greater number of functional synapses^23^, unsilenced synapses^24^ or increased presynaptic release probability^22^ explain the increased frequency. New or unsilenced synapses could increase mEPSC frequency independently of altering presynaptic release probability. BDNF increased mEPSC frequency without changing amplitude while simultaneously increasing both docked vesicle numbers and excitatory synapse density, which mean that the same electrophysiological pattern can arise from concurrent presynaptic and structural changes^23^. Due to the existing evidence on structural neuroplasticity with psychedelics, it is likely that formation of novel spines^5,13^ is reflected functionally as mEPSC frequency increase already 24 h after treatment, potentially also constituting re-wiring of neuronal circuits^12^.

We did not detect psilocybin-induced changes in mEPSC amplitude. This suggests that psilocybin leaves the total of somatically detected quantal currents from individual active synapses largely preserved, not changing AMPA receptor number and conductance^31^. This observation points against general, cell-wide postsynaptic AMPA receptor scaling or an AMPA receptor-mediated synaptic potentiation. Our results are largely line with previous research which has reported that in vivo treatment with psychedelics significantly affects frequency but not amplitude changes of spontaneous excitatory postsynaptic events^5,32–34^. Significant increases of mEPSC amplitudes after 1 mg/kg psilocybin have been observed in the ILA^9^. Arguably, the ILA is the only brain area where we found a trending increase in mEPSC amplitude size. However, mEPSC amplitudes remaining unchanged does not exclude postsynaptic remodeling in a proportion of synapses, as synaptic plasticity may be input-specific^35^ or postsynaptic effects being detectable at a later time-point^36^.

The area-specific kinetic changes showed an additional form of synaptic remodeling that was not captured by the global frequency analysis. In the ACAd, psilocybin shortened mEPSC decay and rise time, whereas in the BLA, it prolonged decay and rise time. Different brain regions may drive temporal integration of psilocybin-modulated excitatory input in distinct ways. A prolonged decay in the BLA is expected to increase charge transfer and temporal summation for an event of a given peak amplitude^37^. Because BLA also showed increased mEPSC frequency, these two changes likely sum up to produce a particularly strong enhancement of integrated excitatory drive. Conversely, faster rise and decay kinetics in ACAd, in the absence of a frequency increase, could narrow the integration window and increase the temporal precision of synaptic signaling^37^. Shorter decay time indicates the presence of AMPA receptors lacking the subunit GluA2^28^, a subunit-composition typically occurring transiently before long-term potentiation manifests^38^. A positive relationship between rise and decay time can reflect dendritic filtering^26^, and structural changes in the synapse and surrounding neuropil can alter the glutamate waveform experienced by AMPA receptors without a change in receptor composition^39^.

Our local *Htr2a* deletion experiment provides evidence for the necessity of 5-HT_2A_ receptors in psilocybin-induced plasticity. Psilocybin reproduced the frequency increase in Rosa26-control neurons but failed to increase mean mEPSC frequency after local *Htr2a* deletion. This finding supports a postsynaptic requirement for 5-HT_2A_ receptor expression for the induction of long-lasting frequency changes in the targeted AI/ORBvl circuit. It is consistent with recent evidence that targeted deletion of 5-HT_2A_ receptors from medial frontal pyramidal neurons abolishes psilocybin-induced spine remodeling and stress-related behavioral effects^13^, as well as identifying intracellular 5-HT_2A_ receptors as important mediators of psychedelic-induced neuronal growth^11^. Postsynaptic re-expression of 5-HT_2A_ receptors is sufficient to rescue DOI-mediated synaptic plasticity^40^, however, presynaptic 5-HT_2A_ receptors on long-range inputs are sufficient to mediate synaptic plasticity and spine formation in cortical areas endogenously not expressing 5-HT_2A_ receptors^33^. Local circuits are also affected by genetic manipulation like our loss-of-function experiment or post-synaptic rescue^40^. Therefore, presynaptic 5-HT_2A_ receptors on short-range inputs in theory could be contributors to synaptic plasticity. In the mPFC, long-range inputs outweigh inputs from local circuits^41^, which may be similar in the AI/ORBvl where we conducted the KO experiment. It is therefore likely that the postsynaptic 5-HT_2A_ receptors are the major contributors to the excitatory plasticity assessed here. While the knockout experiments demonstrated local necessity of 5-HT_2A_ receptors, there was a residual shift in the inter-event-interval distribution curve, which may be explained due to incomplete ablation of the receptor, circuit-level effects from non-transduced neurons, or signaling through additional psilocin targets.

Spatially defined data provide a useful framework for relating regional electrophysiological sensitivity to the distribution of drug-targeted receptors. There is remarkable heterogeneity in 5-HT receptor expression within and between cortical areas on the mRNA level^13,33,42,43^ as well as on the protein level^44^. The correlation between mEPSC frequency and expression of *Htr2a*, *Htr2c*, and *Htr1a* in psilocybin-treated mice substantiate that the cellular and regional expression patterns of psilocin-targeted receptors affect the neuronal response to psilocybin. The 5-HT_2A_-^13^ but also the 5-HT_2C_ receptor^9^ have been implication in psilocybin-mediated neuronal plasticity. Along this line, we found a robust frequency increase despite relatively low *Htr2a* expression in the BLA. In the BLA, plasticity effects may be mediated by upstream afferents or other 5-HT receptor subtypes like the 5-HT_2C_ receptor, which we found highly abundant in the BLA. The psilocybin-specific associations with *Htr2c* and *Htr1a* are also compatible with evidence that 5-HT receptor subtypes other than the 5-HT_2A_ receptor can modulate post-acute behavioral effects. For example, both 5-HT_1A_- and 5-HT_2A_ receptor-related mechanisms contributed to psilocybin-enhanced cognitive flexibility in an activity-based anorexia model^45^. A correlation between *Htr2a* expression and mEPSC frequency was also present in neurons from saline-treatment. This indicates that *Htr2a* expression may reflect pre-existing differences in cortical organization, cellular specialization or baseline excitatory connectivity, rather than solely predicting the magnitude of the psilocybin response. In addition, we observed that *Htr2a* receptor expression and mEPSC frequency showed anterior-posterior gradients, potentially constituting a spatial autocorrelation.

The spatial distribution of the observed effect suggests that psilocybin-induced plasticity preferentially engages limbic circuits including anterior cortical areas. The regions where we observed psilocybin-induced plasticity are critical for value updating, behavioral strategy selection, salience processing, and emotional learning^46^. Given the established involvement of ORBvl in reversal-dependent updating of stimulus-outcome and action-outcome contingencies^47,48^, and involvement of the PL in the encoding and contingency-dependent control of goal-directed actions^49,50^, the enhanced excitatory input observed in these regions may contribute to changes in associative updating after psilocybin treatment. Psilocybin has been reported to enhance set-shifting performance in rats^51^. The regional effects observed in AI and BLA are notable because AI-involved circuits have established roles in processing visceral, salient, and valenced information, whereas the BLA contributes to the updating of affective value and threat-related associations^52–55^. Whether the synaptic changes identified here modify these functions remains to be determined. The observed subregional specificity is notable. ORBvl, but not ORBm, showed a frequency increase, and PL was affected whereas adjacent ILA and ACAd were not. These differences indicate that broad categories such as “prefrontal cortex” may hide different plasticity responses across neighboring subregions. The absence of plasticity in the HPC reported here should not be interpreted as the hippocampus not being sensitive to psilocybin. Psilocybin has strengthened hippocampal excitatory synaptic transmission in chronically stressed mice^21^ and a higher dose administered in conjunction with fear-extinction training increased hippocampal dendritic complexity, spine density, BDNF–mTOR signaling, and neurogenesis-related markers^56^. Differences in dose, stress history, behavioral experience, hippocampal subfield, cell population, and measurement endpoint could explain these divergent findings.

This study is limited in several points. The recordings performed at 24 h post psilocybin treatment is a snapshot in a presumable dynamic evolution of post-acute plasticity and does not inform on the limits of the lasting effects^36^. Measured of mEPSCs without (opto-)genetic circuit refinement reflect synaptic plasticity regardless of input- and output circuits. The mechanisms underlying psilocybin-modulated AMPA receptor kinetics were not determined. Further experiments would be needed to determine the individual contributions of targeted 5-HT receptor subtypes to area-specific plasticity.

With the present work representing a large, spatially defined assessment of psilocybin-induced functional plasticity, we lay a foundation for understanding the brain-wide consequences of psychedelics’ long-term effects. Building on this foundation, resolving specificity of drug-induced plasticity in brain areas and circuits will inform mechanism-based application of psychedelics treatment as well as management of side-effects outlasting acute intoxication, like drug-induced psychosis. Overall, this will significantly support the application of psychedelics in psychiatry.

## Methods

### Animals

Wildtype male and female C57BL/6JRj mice (Janvier Labs, France) 9–26 weeks old were used. Procedures were approved by Cantonal Veterinary Office Basel-Stadt, in accordance with Swiss law. For sacrifice, mice were deeply anesthetized with 5% isoflurane and rapidly decapitated.

### Drug treatments

Psilocybin was obtained as psilocybin*2H_2_O from Apotheke Dr. Hysek AG (Biel, Switzerland). For treatment, psilocybin was dissolved in 0.9% NaCl to 0.1 mg/ml of its free base. Mice were injected i.p. with a single dose of 1 mg/kg psilocybin or an equivalent volume of saline. Treatment allocation was concealed from the involved experimenters throughout drug administration, experiment, and data analysis.

### Head-twitch response

Mice were habituated to experimenter handling in five short sessions separated by 1–3 days. On the test day, mice were allowed to acclimatize to the testing room in their home cage for at least two hours. Then, they were individually tested for head-twitch response to psilocybin. They were placed in a transparent circular cylinder (20 cm diameter), after 15 min quickly removed to inject psilocybin (1 mg/kg i.p.) and then placed back for 45 min. Head-twitches were manually scored from top-down video recordings using the freely available software Boris version 9.7.15 (https://www.boris.unito.it/).

### Viral gene transfer

Intracranial virus injections were conducted with standard stereotactic surgeries during deep anesthesia (induction with 5% isoflurane, maintenance at 1–3% isoflurane). For analgesia, mice received 10 mg/ml carprofen from −1 to 4 days relative to surgery and local analgesia (0.5% lidocaine and 0.25% bupivacaine in saline) 10 min before surgery. Viruses AAV9-hSyn-mCherry-Cre (Viral Vector Facility (VVF) of the Neuroscience Center Zürich (ZNZ), Zürich, Switzerland, identifier v147; iCre for plasmid construction: Addgene #24593) and AAV9-FLEX-SaCas9-U6-sgRosa26 (VVF, ZNZ, Zürich, Switzerland, identifier v664; plasmid (Addgene #159914) received from Dr. Larry Zweifel) or AAV9-FLEX-SaCas9-U6-sgHtr2a (VVF, ZNZ, Zürich, Switzerland; plasmid (Addgene #159908) received from Dr. Larry Zweifel) were combined (final titers were 10^11^ vg/ml for Cre virus, 4 x 10^11^ vg/ml for sgRosa26 and sgHtr2a viruses). The virus mixtures were injected bilaterally at 2.2 A/P, ±1.5 M/L, −2.0 D/V from bregma and dura mater at a volume of 500 nl per hemisphere. After 4 weeks of gene expression, mice underwent drug treatment followed by sacrifice for patch-clamping 24 h after treatment.

### Patch-clamp slice electrophysiology

After sacrifice, brains were quickly removed and placed in ice-cold oxygenated artificial cerebrospinal fluid (aCSF) containing (in mM): NaCl 119, KCl 2.5, MgCl_2_ 1.3, CaCl_2_ 2.5, Na_2_HPO_4_ 1.0, NaHCO_3_ 26.2 and glucose 11. Coronal slices (220 µm thick) were cut with a vibratome and maintained at room temperature. All electrophysiology recordings were performed in a recording chamber perfused at 2.5 ml/min with aCSF at 28 °C. Putative layer 5/6 pyramidal neurons were identified visually under infrared differential interference contrast optics on the basis of their pyramidal somata which are located typically 125–400 µm from the L1/2 border. Patch electrodes were pulled from borosilicate glass capillaries to a tip resistance of 2–4 MΩ. The internal solution was made of (in mM) CsMeSO_3_ 120, CsCl 10, creatine phosphate 5, Na_2_ATP 4, Na_3_GTP 0.4, EGTA 10, HEPES 10 adjusted to pH = 7.3 and 300 mOsm. In some recordings, 0.1% biocytin was included in the internal solution. AMPA receptor-mediated mEPSCs were recorded in the presence of 100 µM picrotoxin and 1 µM tetrodotoxin at a holding potential of −70 mV. Signals were amplified (Multiclamp 700B, Axon Instruments), filtered at 3 kHz and digitized at 10 kHz (Digidata 1550B, Axon Instruments) and collected for 3–5 min using pClamp 11.3 (Molecular Devices) in a gap free protocol 5–15 min after breaking through the cell membrane. Recordings were rejected if access resistance changed by more than 20% or membrane resistance was less than 100 MΩ. The approximate locations of the patched neurons were mapped at the time of recording in the “Mark where you patched” interactive viewer. Recorded traces were processed using Easy Electrophysiology (version 2.4.0). First, traces were low-pass filtered with 1000 Hz Bessel to have better root mean square to analyze. Then, putative events were identified with threshold-based detection (negative peak direction, 5 ms local maximum period, 30 ms decay search period, 5 pA threshold, 10 ms search period, 1-ms averaged baseline, curved baseline and threshold). Mean rise- and decay times were calculated from 200–300 fully isolated mEPSC events per cell.

### The “Marked where you patched” interactive viewer recourse

To document the anatomical location of each recorded cell and archive the CCF coordinates, we developed “Mark where you patched”, a lightweight, browser-based tool for annotating patch-clamp sites on the Allen Mouse Brain Common Coordinate Framework (CCFv3)^25^. The tool displays coronal sections of the 25-µm CCFv3 average template with overlaid region contours. Clicking a location returns the corresponding brain region together with its CCF (µm) and Bregma-referenced (AP/DV/ML, mm) coordinates. Recorded points can be annotated with free-text notes and exported as CSV, JSON, SVG, or image files for downstream analysis and figure preparation.

### Biocytin staining and imaging

After whole-cell patch-clamp recordings, brain slices were fixed in 4% paraformaldehyde in phosphate-buffered saline (PBS) for 24 h at 4 °C. Then, slices were washed three times for 10 min each in PBS and kept in PBS at 4 °C. For staining, slices were washed three times in PBS for 10 min, then blocked and permeabilized for 2 h with PBS containing bovine serum albumine 10% and Triton X 1%. Then, slices were incubated for 72 h at 4 °C with streptavidin conjugated to Alexa Fluor™ 488 (Thermo Fisher Scientific, Cat.# S32354) diluted 1:500 in PBS containing 3% bovine serum albumin and 0.3% Triton X. Then slices were washed three times for 10 min in PBS and incubated for 1 h with 2 µg/ml Hoechst 33258 (Sigma Aldrich, Cat.# 94403) in PBS. Slices were washed and three times for 10 min in PBS, slices were mounted with Vectashield Antifade Mounting Medium. Washing was conducted at room temperature. Images were captured using a Leica SP8 confocal microscope at 20x magnification with z-step size of 0.16 μm.

### Spatial transcriptomics analysis

An open-source spatial transcriptomics data set and associated metadata^19^ were downloaded through AbcProjectCache (https://alleninstitute.github.io/abc_atlas_access/notebooks/abc_atlas_selection_example.html, accessed January 2026). This data set is based on single cell RNA-sequencing results imputed onto multiplexed error-robust fluorescence in situ hybridization (MERFISH) transcriptomics from brain of 7–10 weeks old C57BL/6J mice. A DataFrame with expression of genes of interests (log2(CPM+1)) matrix and metadata of each cell, including “ccf_coordinates”, “neurotransmitter”, and “parcellation_structure”, was retrieved. The data was filtered to restrict for gene expression in glutamatergic cells. To estimate local gene expression profiles for patch-clamp recorded cells, mEPSC data were matched to single-cell spatial transcriptomic datasets based on anatomical region (“parcellation_structure”) and CCF position (“ccf_coordinates”). The transcriptomics data set was loaded and restricted to glutamatergic neurons. mEPSC data were imported from a separate CSV table. Region names in the mEPSC dataset were mapped to the corresponding “parcellation_structure” labels used in the transcriptomic metadata. To spatially align cells across datasets, CCF x coordinates were discretized into bins of width 0.1 mm. For each transcriptomic dataset, cells were assigned to an x-axis bin by flooring the x-CCF coordinate to the nearest lower bin edge. Within each combination of “parcellation_structure” and x-bin, the mean expression of selected target genes was computed, and the number of cells contributing to each bin was recorded. We selected the genes *Htr2a, Htr2c* and *Htr1a* since these are pharmacological targets of psilocin^14,15^, and *Slc17a7* as a control gene. For each cell in the mEPSC dataset, the x-CCF coordinate was assigned to an x-bin using the binning procedure described above. A matching transcriptomic bin was then identified based on the combination of mapped brain region and x-bin and the mean gene expression values from the identified region-bin were assigned to the corresponding mEPSC cell. The number of transcriptomic cells contributing to the matched bin was recorded. In average 690 transcriptomics cells were assigned to a mEPSC cell. Using this procedure, each patch-clamp recorded cell was assigned a region- and position-matched estimate of average local gene expression, which was subsequently exported for correlation analysis.

### Statistical analysis and data visualization

Before two-group comparisons, data were first tested for normality using Shapiro-Wilk test and then either Mann-Whitney or t-tests (with or without Welch correction) were performed. For datasets containing two variables, two-way ANOVA was performed followed by post-hoc multiple comparison testing with false discovery rate (FDR) correction using the Benjamini-Krieger-Yekutieli method when a significant main effect was detected. For correlation analyses, data were fit using linear regression, and correlation strength was assessed using Pearson’s correlation coefficient.

Graphs were generated using the software GraphPad Prism or Python using matplotlib and seaborn. To visualize data in the mouse brain atlas, reference atlas data was obtained from the AllenSDK MouseConnectivityCache at 10 μm resolution, and anatomical boundaries were derived from the Allen CCF annotation volume.

## Data availability

The data will be made available upon publication of this manuscript.

## Code availability

“Mark where you patched” is distributed as a single self-contained HTML file and is freely available at https://github.com/Simmler-lab/Mark-where-you-patched under the MIT license. The underlying atlas data are derived from the Allen Mouse Brain Common Coordinate Framework (CCFv3), © Allen Institute for Brain Science, and are used under the Allen Institute Terms of Use (https://alleninstitute.org/legal/terms-use/).

## Acknowledgements

This work was supported by the Swiss National Science Foundation (SNSF; Grant No. TMSGI3_211261 to LDS). We thank Dr. Larry Zweifel, University of Washington, for providing plasmids for *Htr2a*- and *Rosa26*-KO, and Dr. Jean-Charles Paterna and Dr. Lazaros Vasilikos from the Viral Vector Facility, Neuroscience Center Zürich for virus production.

## Authors contributions

ZL and LDS conceptualized the study. ZL performed patch-clamp experiments and computational analyses of transcriptomic data. FS, CW and ZL performed behavioral experiments. CW and ZL performed immunostaining and microscopy imaging. ZL, LDS, CW and FS analyzed the data. ZL and LDS wrote the manuscript. All authors reviewed and approved the final version.

