## Supplementary Information for "Mapping excitatory synaptic plasticity evoked by single-dose psilocybin in mice"

for

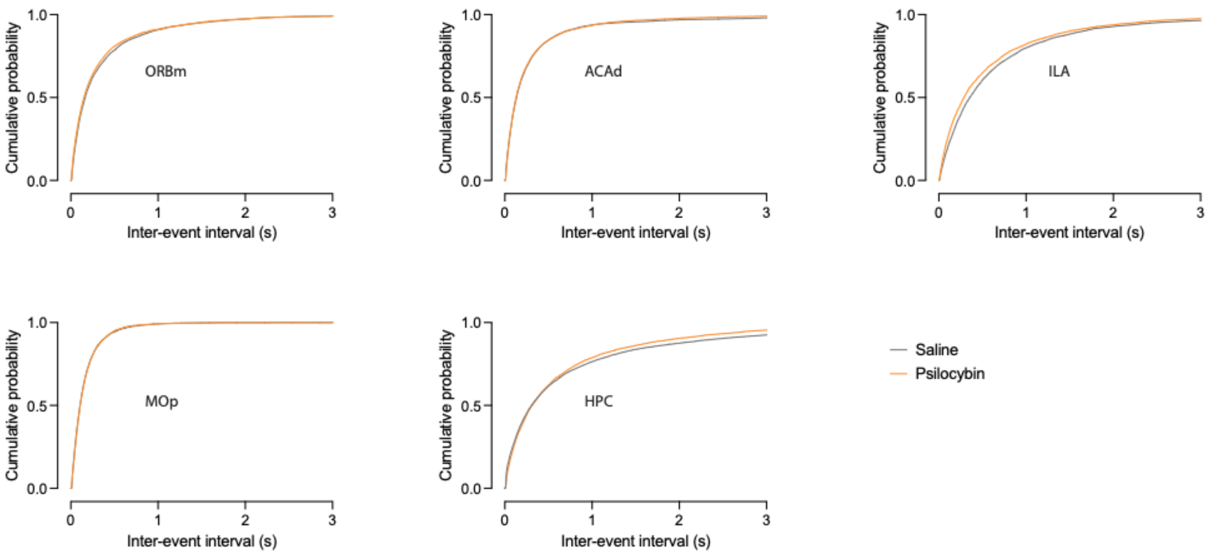


***Figure S1: Cumulative curves for inter-event intervals.***
*Mean cumulative curves for inter-event intervals from mEPSC frequency presented in Figure 2 for ORBm, ACAd, ILA, MOp, and HPC.*
*ORBm, medial orbital area; ACAd dorsal anterior cingulate area; ILA, infralimbic area; MOp, primary motor area; HPC, hippocampus CA1.*

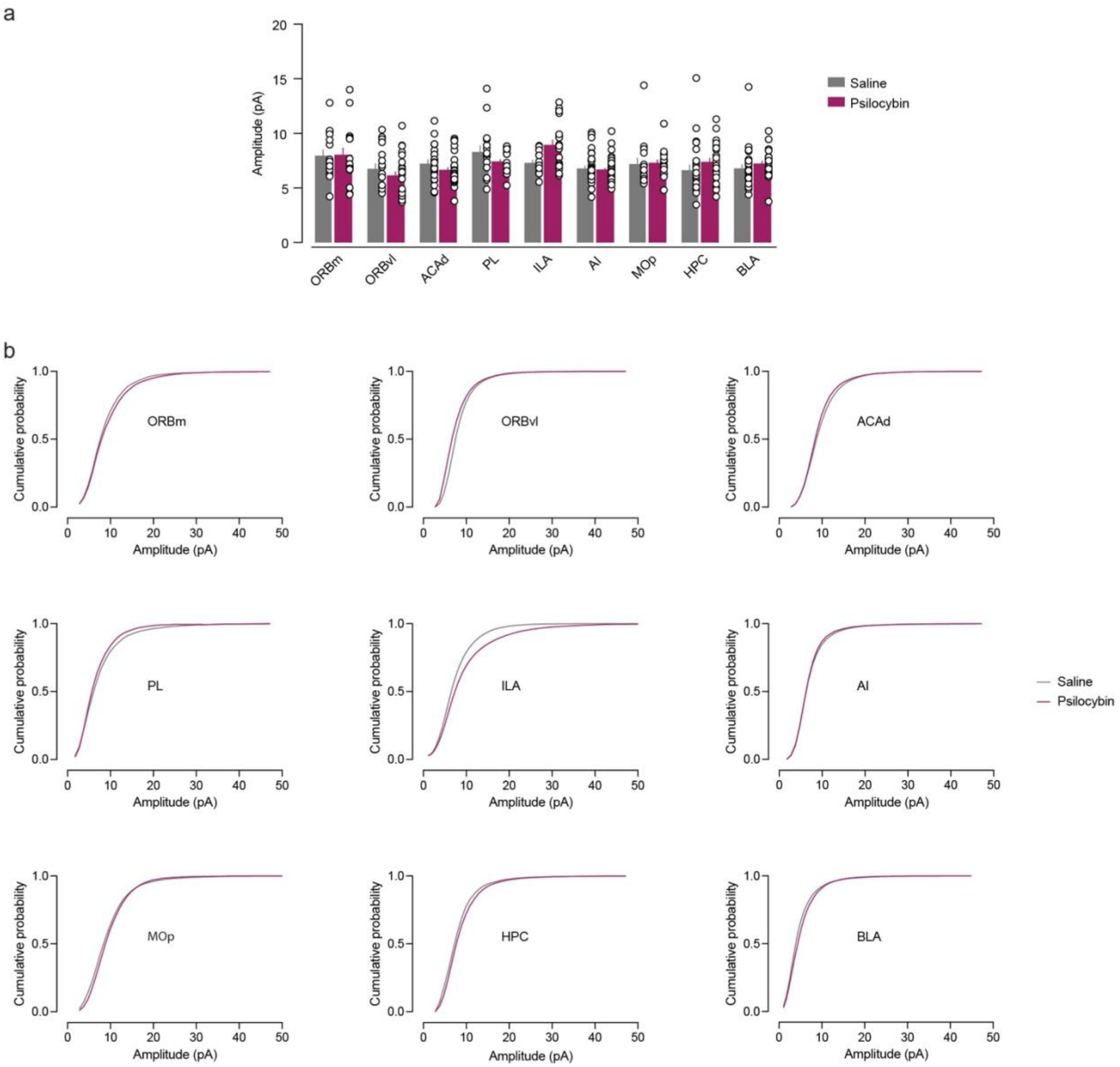


***Figure S2: Amplitudes of mEPSC data in different brain areas.***

***a)*** *mEPSC amplitudes from Figure 1 resolved by targeted brain area. No significant main effect in two-way ANOVA. N(each condition)=16–35 cells from 5–8 mice. Data are mean±SEM with cells as individual data points.*

***b)*** *Data from a displayed as mean cumulative curves.*

*ORBm, medial orbital area; ORBvl, ventrolateral orbital area; ACAd dorsal anterior cingulate area; PL, prelimbic area; ILA, infralimbic area; AI, agranular insular area; MOp, primary motor area; HPC, hippocampus CA1; BLA, basolateral amygdala.*

**Table S1:** Details on mouse and cell numbers in mEPSC data.

| Treatment group | Brain region | Number of mice | Number of cells |
|---|---|---|---|
| Saline | ORBm | 7 | 16 |
| Saline | ORBvl | 5 | 18 |
| Saline | ACAd | 8 | 22 |
| Saline | PrL | 6 | 16 |
| Saline | ILA | 7 | 17 |
| Saline | AI | 5 | 33 |
| Saline | MOp | 6 | 16 |
| Saline | HPC | 6 | 26 |
| Saline | BLA | 8 | 28 |
| **Total saline*** | | **34** | **192** |
| Psilocybin | ORBm | 7 | 16 |
| Psilocybin | ORBvl | 5 | 23 |
| Psilocybin | ACAd | 7 | 29 |
| Psilocybin | PrL | 7 | 21 |
| Psilocybin | ILA | 6 | 18 |
| Psilocybin | AI | 6 | 35 |
| Psilocybin | MOp | 6 | 16 |
| Psilocybin | HPC | 5 | 21 |
| Psilocybin | BLA | 6 | 21 |
| **Total psilocybin*** | | **35** | **200** |

*Numbers do not add up because for several mice, cells in more than one brain region were recorded.
